# Disentangling Cephalopod Chromatophores Motor Units with Computer Vision

**DOI:** 10.64898/2025.11.30.691401

**Authors:** Mathieu D. M. Renard, Johann Ukrow, Margot Elmaleh, Dominic A. Evans, Yifan Wu, Xitong Liang, Gilles Laurent

**Affiliations:** Max Planck Institute for Brain Research, Frankfurt, Germany; State Key Laboratory of Membrane Biology, School of Life Sciences, Peking University, Beijing 100871, China; IDG/McGovern Institute for Brain Research, Peking University, Beijing 100871, China; Peking-Tsinghua Center for Life Sciences, Academy for Advanced Interdisciplinary Studies, Peking University, Beijing 100871, China; Center for Quantitative Biology, Academy for Advanced Interdisciplinary Studies, Peking University, Beijing 100871, China

## Abstract

Cephalopod chromatophores are skin pigment organs that enable unmatched camouflage through rapid, flexible and neurally controlled deformation. Although their morphology is well known, the organization of their motor control is not entirely understood. Here, we combine high-resolution videography with a dedicated computer-vision pipeline (CHROMAS) to investigate chromatophore control and their likely innervation in *Euprymna berryi* and *Sepia officinalis*. By segmenting chromatophores into radial slices and analyzing anisotropic deformations, we applied dimensionality reduction (PCA) and source separation (ICA) to estimate the number and spatial influence of motor neurons responsible for the control of individual and groups of chromatophores. On average, four independent components were detected (suggesting innervation by four motor neurons), each forming contiguous petal-shaped domains rather than causing uniform expansion. Clustering thousands of components revealed motor units spanning multiple chromatophores, most involving fewer than 14 but occasionally spanning more widely. These motor units displayed a wide variety of geometries, ranging from compact local groups to elongated or fragmented structures; they often overlapped, with repeated co-innervation of chromatophore pairs occurring more often than expected by chance. Expansion was consistently faster and more stereotyped than relaxation, consistent with active contraction (corresponding to chromatophore expansion) and passive recoil (chromatophore contraction). Together, these results show that individual chromatophores are not singular or uniform pixels, but rather contrast elements that can be fractionated into smaller territories, themselves coordinated with those of other chromatophores. This geometry of neural control enables, among others, the generation of “virtual” chromatophores, i.e., functional groupings of adjacent chromatophore territories that act as single units, as well as that of noise in the distribution of pixel shapes.

## Introduction

Coleoids, a subgroup of the cephalopod taxon which includes octopuses, squids, and cuttlefish, possess one of the most sophisticated camouflage systems found in nature. This extraordinary ability is mediated by arrays of pigment cells, known as chromatophores, which expand and contract to alter skin coloration and statistical patterning in response to motor command from the brain. The collective behavior of thousands or millions of chromatophores thus creates adaptive patterns that enable the animals to blend with their surroundings, evade detection by predators or prey and communicate with conspecifics (Hanlon & Messenger, 1988).

Cephalopod chromatophores differ from pigment cells in other animals: each chromatophore is a neuromuscular organ composed of a central pigment cell containing a cytoelastic sacculus, surrounded by 15–25 slender radial muscle fibers, themselves attached to an extracellular semi-stiff fiber mesh at their distal end (Messenger, 2001). The pigment granules residing in the elastic sacculus thus spread into a thin disc when the radial muscles contract due to motor neuron excitation, expanding the pigmented area. When the motor neuron input ceases, the chromatophore muscles relax, the elasticity of the sacculus causes it to recoil (and the cytoplasmic membrane to invaginate and fold onto itself), reducing visible coloration (or the size of this biological, expandable pixel). These muscle-driven, active expansion and elastic, passive retraction enable very rapid patterning and color changes, seen in no other biological coloration system. In addition to their unique neuromuscular structure, chromatophores occur in distinct color classes (typically yellow, red, brown, and black), each defined by the pigment (e.g, xanthommatin for yellow) and density of pigment granules they contain. These classes are linked to the age of the chromatophore (starting yellow and darkening over several weeks; Reiter et al., 2018), and usually migrate through the thickness of the skin, thus forming superimposed chromatic layers. Together they broaden the chromatic range available for patterning.

The basic neural circuit for camouflage patterning is well established: motor commands flow from visual processing centers (optic lobes) through basal lobes to the chromatophore lobes, where the somata of the motor neurons that directly innervate the muscles of the skin chromatophores reside (Boycott, 1961; Messenger, 2001). What remains unclear is how the axonal terminals of these motor neurons map onto individual and groups of chromatophores. Anatomical and physiological studies from Florey and colleagues (Cloney & Florey, 1968; Florey, 1969), as well as Dubas and Boyle (1985), provided evidence for polyneuronal innervation, with individual muscles or chromatophores under the control of several motor neurons. However, the number of inputs, their spatial organization, and the logic of motor unit formation remain unresolved.

This uncertainty limits our understanding of how the distributed neural activity that controls camouflage generates the details of body patterns. Treating chromatophores as uniform “pixels” ignores the possibility that subregions within each chromatophore could act as semi-independent effectors. Likewise, the degree to which chromatophores are coordinated in groups—forming motor units that span several chromatophores—has only been estimated qualitatively (Florey, 1969; Dubas & Boyle, 1985). These early anatomical and physiological studies suggested that individual motor neurons may innervate dozens to hundreds of chromatophores, but such estimates were indirect and limited in scope. More recently, computational approaches have begun to address this issue (Reiter et al., 2018), but these analyses considered chromatophores as indivisible units and provided only indirect evidence of motor-unit structure. No systematic, quantitative framework yet exists to describe the number, geometry, and overlap of chromatophore motor units on a large scale. We addressed this gap using high-resolution tracking of chromatophore dynamics combined with computational tools capable of disentangling overlapping motor inputs.

## Results

### Methodological development with Euprymna berryi

To establish a quantitative framework to analyze chromatophore motor control, we first focused on a model system with easily resolvable chromatophores. We chose the hummingbird bobtail squid, *E.*D*berryi*, because of large chromatophores, compact size, and rapid growth. We first show the analysis of a 30-s video dataset (1080 × 1080 px, 20 frames per second or fps) taken from a sedated hatchling (Fig. 1a; see Methods). Because our primary aim was to describe the composition and coordination of chromatophore motor units, it was important to examine animals in the absence of the descending commands that occur during active behavior. Spontaneous activity, typically mild and “noisy” was thus ideal to enable measurements of the motion correlations between chromatophores that reflected shared motor neuron drive, rather than shared correlations due to upstream motor neuron groupings by premotor circuits.

Chromatophores were segmented, each divided in polar (radial) slices and the areal variations of each slice across all successive time frames were measured, as developed and described in Ukrow, Renard et al., (2025). Chromatophore slices were then clustered using HDBSCAN (Hierarchical Density-Based Spatial Clustering of Applications with Noise, Campello et al., 2013), based on the temporal correlation of their size variations. This analysis revealed that clusters were composed of slices that lay both within and across individual chromatophores (Fig. 1b-c), with some chromatophores contributing to multiple clusters (Fig. 1d-f). Within a cluster, the orientation of each slice tended to align centripetally towards a virtual point located between the contributing chromatophores (Fig. 1g), effectively forming a closed “virtual chromatophore” rather than ring-like patterns. By “virtual chromatophore” we thus refer to a functional unit defined by the coordinated slice activities of adjacent chromatophores that share motoneuron innervation. Notably, smaller (yellow, less mature) chromatophores in the same region did not deform and therefore did not cluster, despite lying in the same apparent innervation fields. These inactive chromatophores proved to be very useful as stable anchors for motion tracking. If *E. berryi* allowed us to identify the important parameters of our analyses, this nocturnal species is only moderately interesting from a camouflage point of view. We thus transferred our methods to *Sepia officinalis*, a diurnal master of camouflage with large numbers and high density of chromatophores.

### Analysis of chromatophore motion in Sepia officinalis

We applied CHROMAS to the skin of *S. officinalis*, where chromatophore density is roughly ten times higher and chromatophore dynamics are correspondingly harder to resolve. This animal (Fig. 2) was recorded under head fixation with and without sedation (HF+S and HF), conditions chosen to elicit spontaneous chromatophore activity required for correlation analysis (see above), using 8K video recordings and visible implant elastomer (VIE) markers for consistent region identification across days or weeks (see Methods). We focused on a small region at the center of the dorsal mantle (Fig. S1) and analyzed sections (0.5 by 0.5 mm) of a 108-s HF video segment (2160 images) and a 20-s HF+S segment (400 images), that together yielded 3,285 segmented individual chromatophores. These datasets provide the basis for the following quantification of chromatophore anisotropic expansion and motor unit identification.

### Interpretation of fine single-chromatophore motion

We performed PCA on the 36Lslice time series of each chromatophore (see Fig. 2a for a single chromatophore) and from it estimated the number of principal components (PCs) needed to explain most of the variance in the motion data. The number of relevant components was determined by identifying the point of maximum curvature in the cumulative explained variance curve, as implemented in the CHROMAS pipeline (Ukrow, Renard et al., 2025; see Methods). In Fig. 2b, this point corresponds to PC5 (red dashed line), beyond which additional components contribute negligibly to the explained variance. Over 1,829 chromatophores in the HF□+□S dataset, this approach yielded 3.715 ± 1.211 PCs per chromatophore on average (Fig. 2c). To test for an effect of sedation, we repeated the analysis on 1,456 chromatophores in the head-fixed, not sedated condition (HF), and obtained 3.683 ± 1.224 PCs. A Welch’s t-test reveals no significant difference between these two conditions (t = 0.21, p = 0.83; Cohen’s d = –0.03).

The number of dominant explanatory components identified by PCA was then used to initialise an Independent Component Analysis (ICA). This operation extracted the dynamics of the putative underlying sources (Fig. 2d) and revealed their spatial influence on individual chromatophore slices (Fig. 2e–f). Each independent component—which we interpret as the signature of firing activity in a single motor neuron—was overlaid onto the 36 equal polar slices to generate a detailed influence map for each chromatophore (Fig. 2e-f). Across our dataset, the putative motor neurons, so identified, innervated between three and sixteen slices per chromatophore. Importantly, these targeted slices consistently formed contiguous domains around the chromatophore’s circumference. The influence profiles frequently adopted a petal-like shape, characterised by a peak in one polar position and a gradual tapering of influence across the adjacent slices. This parcelated representation suggests the topography of motor innervation of single chromatophores, and provides the most likely explanation for the observed anisotropic expansion patterns of single chromatophores.

### Structure and size of chromatophore motor units

Affinity-propagation clustering of the 6,795 independent components (ICs) extracted from 1,829 chromatophores (HF+S dataset) yielded 754 clusters, and thus putative motor units (MUs). Each putative MU comprised a mean of 9.003 ± 4.755 ICs, and 95.877 ± 7.566 % of these MUs spanned multiple chromatophores, indicating extensive multi-chromatophore innervation (Fig. 3a-b). For this spatially bounded dataset (Fig. 3b), the cluster-size distribution was skewed, with 90 % of motor units innervating fewer than 14 chromatophores (Fig. 3c). We analyzed the same chromatophores under our two conditions: HF and HF+S. A Welch’s t-test revealed no significant difference in the number of components per MU between the two groups (HF = 8.551 ± 5.753, n = 626, vs. HF+S = 9.003 ± 4.755, n = 754; t ≈ –1.57, p = 0.12).

The spatial relationship between chromatophores belonging to the same putative MU was assessed by calculating pairwise distances and convex hull areas of putative MU clusters. The global nearest-neighbor distance (NND)—defined as the average shortest distance from each chromatophore to its closest neighbor across the entire dataset—was 6.130L±L1.589Lµm. In contrast, the mean within-cluster NND was 40.548□±□46.946Lµm, indicating that chromatophores grouped within the same motor unit are not limited to immediate neighboring chromatophores. The corresponding furthest distance—defined as the greatest distance between any two chromatophores within a cluster—was 228.96□±□94.36□µm. The corresponding global value (361.50□±□49.80Lµm) depends directly on the field of view of the camera and is thus less informative.

To quantify spatial coverage, the convex hull area was measured for each cluster. (The convex hull area refers to the area of the smallest convex polygon that fully encloses a given set of points.) The distribution of these areas (Fig. 3d) shows that most clusters occupy relatively small regions. Across 754 clusters, convex hull areas ranged from 25.18□µm^2^ to 915,087.66□µm^2^, with a median of 136,199.10□µm^2^ and a mean of 209,793.70□µm^2^. Ninety percent of clusters had a convex hull area smaller than 51,513.25 µm^2^ (or 0.0515 mm^2^, the equivalent of a fine grain of sand). Figure 3e-f illustrates the relationships between spatial and compositional features of motor units. A positive correlation (*r*□=L0.66) was observed between cluster area and chromatophore count (Fig. 3e), while a weak negative correlation was found between area and chromatophore density (*r*□=□–0.14) (Fig. 3f), and between chromatophore count and density (*r*□=□–0.14). These relationships suggest that clusters occupying larger areas tend to have more chromatophores (Fig. 3e), but at lower density (Fig. 3f).

We next examined the secondLorder coLoccurrence, defined as the number of unique chromatophore pairs that were parts of at least two distinct motor units. This number was 1,073 in our dataset and was compared to the value expected under a null hypothesis of random paired assignments. A set of 1,000 random permutations yielded a distribution shown in Fig. 3g, and an expected mean value of 263.72 ± 16.94 pairings. The highly significant difference between these two values (Z = 49.36, p < 0.0001) indicates that co-innervations are not random, and suggests the existence of cooperative or joint-causality mechanisms for the innervation and control of nearby chromatophores.

The spatial structure of putative motor units displayed a wide range of configurations (Fig. 4). Some clusters were compact, with chromatophores arranged closely together and spanning small distances (e.g., panels a–d). In many cases, clusters (i.e., MUs) shared chromatophores, resulting in overlapping network structures (e.g., panels e–f). Other clusters exhibited more elongated or linear profiles (e.g., panel g), with chromatophores extending over a longer axis. In addition, mixed profiles were observed, where a core group of closely spaced chromatophores was connected to one or several distant chromatophore(s) (e.g., panels g and h). Because our field of view did not encompass the mantle in its entirety, our analysis will have missed large motor units spread over very large areas, if they exist.

### Dynamics of chromatophore expansion and contraction

Because chromatophore deformations are caused by active expansion (radial muscle contraction) and passive (elastic) recovery (radial muscle relaxation), the two phases of chromatophore deformation have different dynamics (fast expansion, slower recovery). We tested whether these kinetic differences could be detected in our videographic measurements at 20 fps, i.e., at sampling rates low compared with those of electrophysiology (Florey, 1969). We analyzed expansion and recovery processes on individual slices, so as to isolate the activity of different presumed radial-muscle groups within each chromatophore. We focused on chromatophores with high activity levels to ensure clear detection of expansion and contraction phases and accurate estimation of their respective speeds and durations.

Although very variable, our data showed that chromatophore expansion is significantly faster and more stereotyped than contraction across chromatophores (Fig. 5 and S2). By separating events by amplitude, our videographic approach captured expansion–contraction dynamics across a wide range of events. These results indicate that these kinetic features can be detected with the temporal resolution of our video-imaging. Specifically, across a subset of 10 active chromatophores (from the HF+S dataset), expansion phases occurred at an average speed of 0.667 ± 0.590 pixels/frame, compared with 0.422 ± 0.557 pixels/frame for the contraction (n=2157 events). This difference was significant (paired t-test, t = 19.50, p < 0.001). In our datasets, 10 aligned pixels span 1 μm. Expansion events were spread on average over 4.41 ± 1.91 frames vs. contractions, 5.90 ± 3.50 frames. These results are thus consistent with the basic mechanisms of chromatophore motor activation, and highlight our technique as a reliable tool to study certain aspects of radial muscle biomechanics.

### Direct activation of single chromatophore motor neurons (Euprymna berryi)

The above observations in *Sepia* were based on high-resolution video image data of spontaneous resting activity, enabling the simultaneous monitoring of large regions of skin and thus, the simultaneous tracking of large numbers of chromatophores and measurements of their motion correlations. To test for the validity of the approach, we also carried out combined video imaging and whole-cell patch-clamp recordings of single motor neurons, thus enabling the direct control of single-motor neuron discharge. Our goal was to test, on a small scale, the validity of the large-scale motion-correlations approach based on measurements of spontaneous activity. In particular, we wished to check directly whether individual chromatophores are anisotropically controlled by multiple motor neurons with polarised action (as suggested by the deformation “petals” in Fig. 2), and whether the action of individual motor neurons on a field of nearby chromatophores is usually directional, with expansion vectors oriented towards a virtual center of mass.

To this end, we returned to *Euprymna* (to increase the probability of finding both motor neurons and corresponding chromatophores, because the number of chromatophores in this species is small), patched a motor neuron in the posterior chromatophore lobe and looked for the patch of skin containing the innervated chromatophores. Once those had been identified, we injected a 1s-long direct current in the patched motor neuron to cause sustained firing (Fig. 6a), and observed the effect of this activation on the chromatophores (Fig. 6b and Fig. SV1). Here the motor neuron innervates five nearby chromatophores, causing a partial, directed, petal-like expansion of each one towards a virtual common center located near their common center of mass. Detecting this innervation pattern in this example proved difficult using spontaneous activity alone (*i.e.*, excluding the electrophysiological recording), due to the small amplitude of the chromatophore deformations caused by single motor neuron action potentials. These observations were repeated in 16 motor neurons from 12 different animals, revealing a detectable effect in 4.38 ± 0.55 chromatophores (mean ± SEM, n = 16) (Fig. 6c; videos provided in the Data Availability section). Note however that the expansion of chromatophores by single-motor neuron electrical activation was not always anisotropic (Fig. 6c - bottom right), suggesting that some chromatophores in *Euprymna* are controlled by single motor neurons (or that motor neurons with a common chromatophore target may be electrically coupled). We conclude that our quantitative videographic approach based on the analysis of spontaneous chromatophore deformations in *Sepia* reveals features that are consistent, but on a large scale, with those revealed by focal motor stimulation.

## Discussion

Our results extend early focal electrophysiological observations to large physical scales, thanks to the quantitative and easy-to-use videographic methods that we developed recently (Ukrow, Renard et al., 2025). Messenger (2001), drawing on Florey’s electrophysiological studies (Florey, 1969), suggested that chromatophores are controlled by at least four motor neurons. Our decomposition consistently revealed three to four independent causes for individual chromatophore deformation. Our results match and support these early inferences, but are here based on direct analysis of thousands of chromatophores in parallel. Likewise, Dubas (1985, 1986) estimated that a single motor neuron may innervate tens to hundreds of chromatophores, typically of the same color. Our clustering revealed units of variable size, typically around ten chromatophores in *S. officinalis* but sometimes extending much further (up to thirty), matching her conclusion that motor fields are both distributed and overlapping. We never detected, however, motor units containing many tens to hundreds of chromatophores with our technique. This may be due to the generally small expansions caused by spontaneous motor neuron discharge, presumably usually limited to single action potentials; it is possible also that the gain of motor neuron-chromatophore transfer varies over the spatial extent of an innervation field. If so, only some of the targets of a motor neuron would be identifiable with our methods.

Contrary to mature skeletal muscle innervation in vertebrates (Henneman, 1957; Kernell, 2006), individual invertebrate muscle fibers often receive convergent excitatory inputs from more than one motor neuron (Hoyle, 1955; Bullock & Horridge, 1965; Sasaki & Burrows, 1998, Florey, 1969; Dubas & Boyle, 1985). The nature of our videographic methods was not sufficient to resolve chromatophore fiber poly-innervation, if it exists. If anything, however, the adjacent deformation “petals” we identified rather suggest adjacent, but non overlapping territories of muscle-fiber innervation by individual excitatory motor neurons. The patterns of correlated deformation of multiple chromatophores were consistent with earlier findings (Dubas & Boyle, 1985; Reiter et al., 2018) that individual motor neurons innervate chromatophores of the same colour. Given the systematic intercalation of yellow and brown chromatophores in the skin (in which chromatophore color is correlated with chromatophore age; Reiter et al., 2018), the monochromatic innervation patterns, combined with their great spatial precision, indicate a remarkable degree of local control of chromatophore innervation by motor neurons in *Sepia*.

Most strikingly, our results reveal a phase offset between the anatomical matrix of chromatophores and their functional innervation. Each chromatophore can be subdivided into several (typically three to four) independently controlled zones, each containing an adjacent subset of the 15–25 radial muscles that control each chromatophore; these zones typically cut across neighboring chromatophores. This results in the possible generation of “virtual chromatophores”, or chromatic zones formed by the convergence of the quadrants of nearby chromatophores. This suggests that the smallest addressable units of cephalopod skin are not necessarily single chromatophores but rather topographically ordered subsets of radial muscles, spread over multiple chromatophores and grouped by their shared motor neuron. This view modifies and extends Packard’s (1982) billboard analogy: chromatophores are not isolated “billboard lightbulbs” but overlapping zones wired through multiple control lines. Such an arrangement implies strong developmental constraints on wiring while simultaneously enhancing the flexibility of body-pattern generation.

Our analyses suggest that chromatophore control operates through overlapping innervation fields that often do not coincide with the anatomical boundaries of individual chromatophores. In *E. berryi*, motor units tended to be compact and converged toward points between neighboring chromatophores, effectively creating “virtual chromatophores” composed of partial contributions from adjacent cells. Several mechanisms could underlie this organization. Developmentally, axons may innervate radial muscles opportunistically within restricted skin regions rather than selectively targeting individual chromatophores, leading to fields that span across cells. The inactivity of younger orange chromatophores embedded within active fields and the independent control of yellow chromatophores suggest that innervation may also be stratified by depth, consistent with the layered organization of chromatophore colors (Messenger, 2001) and the developmental processes shaping this system (Packard, 1982). As chromatophores age and change color, vertical migration within the skin might bring them within other motor innervation territories and lead to their reinnervation. Our techniques will now enable the study of the development of chromatophore innervation and possibly, of their reinnervation as they age and change pigment.

Our videographic approach also provides sufficient temporal resolution to resolve some aspects of radial-muscle kinetics, opening avenues to study chromatophore biomechanics and, ultimately, to separate the contributions of intertwined forces, including but not limited to radial muscles, elastic pigment sacs, and intercellular coupling, to expansion dynamics.

### Functional implications

Although motor units are often composed of multiple chromatophore sectors rather than whole chromatophores, the resulting pigment fields can be visually indistinguishable from those produced by full-chromatophore control. This fragmented organization could simply reflect wiring constraints, but it may also confer functional advantages. By recruiting sectors from different chromatophores, motor units introduce irregular contours, asymmetries, and non-uniform shapes, expanding the expressive range of the skin and reducing geometric repetitiveness in pattern textures. Sector-based units may therefore permit finer spatial modulation or realistic pattern “noise”, particularly during partial activation states.

Most motor units we observed fell below 51,513 µm^2^ in convex-hull area, roughly equivalent to a ∼227 µm square. This scale matches the grain size of very fine sand (125–250 µm; Wentworth, 1922; Blott & Pye, 2012), which dominates Mediterranean coastal habitats where *S. officinalis* is common (Sandulli et al., 2010; Bettoso et al., 2016; Dauvin et al., 2017; Asensio-Montesinos et al., 2022). Such correspondence suggests that motor-unit resolution may be ecologically tuned to substrate textures, providing effective camouflage without incurring the energetic costs of finer-grained control.

Shared innervation of chromatophore sectors by multiple motor neurons may enable gradual, fluid transitions between patterns. As one unit ramps down and another ramps up, overlapping territories smoothen otherwise abrupt boundaries. These overlaps could also underlie dynamic displays such as the “passing cloud” (How et al., 2017; Laan et al., 2014; Mather & Mather, 2003), in which waves of activity propagate seamlessly across the skin.

Motor units exhibited diverse geometries, from compact clusters to elongated or fragmented ensembles. Such variability may be functionally specialized: tight units supporting sharp edges or high-contrast motifs, while dispersed units contribute to gradients or subtle textures. This architecture resembles receptive-field mosaics in sensory systems. Just as retinal ganglion cells (Masland, 2012) or cutaneous mechanoreceptors (Mountcastle, 2005) vary in size and density to capture distinct features, chromatophore motor units form an outward-facing mosaic of overlapping “projective fields” that compose complex visual patterns.

### Technical issues

Our non-invasive recordings and large-scale analyses rely on indirect inference based on spontaneous correlated motion rather than direct electrophysiological measurements. The great advantage of this approach is that it enables large-scale measurements in freely behaving animals. This would be impossible with electrophysiological recordings. Chromatophore expansion served as a proxy for radial muscle contraction, itself a result of motor neuron activity. One limitation is that the inferences we make about motor neuron activation are indirect. Using simultaneous motor neuron recordings and chromatophore imaging in reduced preparations of the brain and mantle skin, we could nevertheless confirm several of our interpretations using direct motor neuron activation. The non-isotropic action of individual motor neurons on individual chromatophores and the converging expansion of nearby chromatophores under the action of a shared motor neuron, for example, could each be confirmed directly. We observed, however, that the direct-current injections needed to generate a detectable chromatophore expansion were often prolonged, causing the production of trains of action potentials in the patched neuron. Spontaneous chromatophore expansion, however, is probably caused by spontaneous and isolated (i.e., single) motor neuron action potentials. It is possible, therefore, that what can be detected using our image analysis approach on spontaneous activity underestimates the true patterns of innervation. This will be especially true if the gains of the many neuromuscular junctions made by a single motor neuron vary across its output field. Only the stronger outputs will lead to detectable motion. (Note that the petal shape of the chromatophore expansion quadrants suggests a gradient of innervation strength—central peak flanked by decreasing motion amplitude over each corresponding quadrant.) We also observe that, to be interpretable, the patterns of chromatophore expansion during spontaneous “noisy” activity should be as decorrelated as possible. If for example, two motor neurons share a presynaptic excitatory drive, they may often fire together when their common presynaptic neuron is active, leading to the erroneous delineation of motor units. Our approach must therefore be applied in conditions of minimal activity, ideally during the apparent spontaneous flicker that is usually observed in resting animals.

Chromatophore slice dynamics were estimated from membrane deformation rather than direct detection of muscle insertion points; similarly, putative motor units were defined by clustering of correlated activity profiles. While such proxy-based approaches involve multiple inferential steps, they are routinely used in neuroscience and physiology to extract functional information from imaging data—for example, inferring spiking from calcium fluorescence or estimating muscle activation from high-speed video recordings. Our strategy applies the same principle to chromatophore dynamics, providing scalable, non-invasive access to living animals.

The two principal clustering strategies which we settled on and used—HDBSCAN, with *E. berryi* during pipeline development, and Affinity Propagation, adopted as a refinement for *S. officinalis*—make different assumptions about cluster structure. In particular, Affinity Propagation groups sectors based on pairwise similarity of activity dynamics rather than local density in feature space. Although both approaches yielded consistent biological insights, it should be noted that inferred motor units are not insensitive to algorithmic choices and should be interpreted with this in mind. More generally, clustering constitutes a form of model selection, and alternative algorithms may reveal complementary aspects of motor-unit organization.

Just as in any imaging approach, spatial and temporal sampling considerations represent important constraints. In *S. officinalis*, recordings were restricted to small regions of the dorsal mantle, precluding whole-body comparisons and leaving open whether motor-unit organization is conserved across skin regions. Developmental factors further narrowed the available time windows for observation: chromatophore overlap in growing *E. berryi* hatchlings and low spontaneous activity in young *S. officinalis* juveniles restricted the stages suitable for analysis. In this context, the large, spatially dispersed motor units observed in *S. officinalis* may not represent a universal architecture; they could reflect the developmental stage of the individuals we studied or species-specific ecological requirements. Previous reports based on more limited, but electrophysiological stimulation experiments, suggest considerable diversity across cephalopods, with Dubas (1985) describing widely distributed motor units in octopus and Florey (1969) reporting more compact units in squid. The extent to which motor unit size and shape vary as functions of species, position on the body, and developmental stage, will need to be examined.

The complex mechanics and innervation patterns of chromatophores introduce ambiguities. Elastic recoil of the pigment sac and mechanical coupling between adjacent radial muscles likely propagate forces beyond the site of neural activation, producing secondary deformations (of unknown amplitude) in neighboring slices. In addition, overlapping motor neuron axons could converge onto the same set of radial muscles, in which case a single slice would at times reflect the combined drive of multiple neurons. Together, these factors make it difficult to determine where the influence of one motor neuron ends and that of another begins. As a result, our estimates should be viewed as functional partitions of activity rather than definitive anatomical boundaries.

In conclusion, we developed an image analysis pipeline (Ukrow, Renard et al., 2025) and used it to analyze the structure of chromatophore expansion and shape control in two species of coleoid cephalopods, from high resolution movies of freely moving animals. This approach revealed many fine features at the distal end of this highly specialized motor system. By identifying first the motor units, as done here, it may become possible to reveal important aspects of their presynaptic drive and thus make precise and testable predictions about the organization of this remarkable neural system.

## Methods

### Chromatophore video acquisition and analysis

#### Animals

All research and animal care procedures related to video acquisition of chromatophores and analysis were carried out in Frankfurt, in accordance with the institutional guidelines that are in compliance with national and international laws and policies (DIRECTIVE 2010/63/EU; German animal welfare act; FELASA guidelines). The study was approved by the appropriate animal welfare authority (Regierungspräsidium Darmstadt) under approval number V54-19c20/15-F126/2012.

Hummingbird bobtail squids *Euprymna berryi* and European cuttlefish *Sepia officinalis* were hatched from eggs laid in house and reared in a seawater system. The closed system contained artificial seawater (ASW; Instant Ocean) with a salinity of 33-36 ppt and pH of 8.1-8.4. Water quality was monitored continuously and was tested regularly. The housing temperature was different in function of the species. Aqua Medic *Tri Complex* supplied macro- and trace elements continuously. A constant water through-flow resulted in 5 complete water exchanges per hour. Room illumination provided a 12□h–12□h light–dark light cycle with gradual on- and off-sets at 07:00 and 19:00. Enrichment consisted of natural fine-grained sand substrate, artificial plastic plants and translucent red plastic houses. Both species were fed live food (either *Neomysis* spp. or small *Palaemonetes* spp.) twice per day.

*E. berryi* were reared in a seawater system at 20-26□°C. The filtration systems included filter bags, protein skimmer, biofilter, UVC-sterilisator and heater. The adult animal groups were housed in □300-L plastic tanks, juveniles in plastic boxes of 18-36 L. Experimental animals of unknown sex, just a few days after hatching, ranging from 3 to 6□mm in mantle length, were selected for healthy appearance and calm behavior.

*S. officinalis* were reared at 18-20□°C. The filtration systems included drum filter, protein skimmer, biofilter, UVC-sterilisator and chiller. The animals were housed individually in 30-90□L PVC tanks with a constant water through-flow resulting in 5 complete water exchanges per hour. Animals above 50 mm in ML were fed once per day defrosted fish or shrimp. Experimental animals of unknown sex, 5 days to 10 months after hatching, ranging from 5 to 100□mm in mantle length, were selected for healthy appearance and calm behavior.

### Arena and filming equipment

*E. berryi:* the arena design aimed to confine the hatchlings within an area fully covered by the camera’s field of view. The resulting arena measured 10 cm × 10 cm × 3 cm in height and included a conical sub-arena with a direct water inflow (Fig. S3a). This cone featured small holes allowing water to outflow into a larger surrounding chamber. The design incorporated a glass lid to reduce visual distortion by enabling direct water-glass contact and improving image clarity. Additionally, the arena floor included a glass surface to permit imaging from below. A peripheral “trench” area surrounding the chamber directed excess water through outflow tubing, ensuring continuous circulation. For recording, we used a Basler Ace 2 (a2A2590-60ucPRO, Basler, Ahrensburg, Germany) equipped with a Kowa lens (LM25JC10M, Kowa, Nagoya, Japan). The imaging configuration enabled 1920 × 1920 px recordings corresponding to a 1 × 1 cm field of view, large enough to encompass the whole body of *E. berryi*. Illumination was provided by a custom-built LED ring mounted above the imaging arena and controlled through a custom software to modulate light intensity cyclically during recordings. Recordings were performed inside a custom sound- and light-proof enclosure lined with acoustic foam to dampen external vibrations and reflections. The box contained the imaging arena, camera, and LED ring, ensuring stable sensory conditions during recordings.

*S. officinalis*: the arena measured 20 cm x 30 cm x 10 cm, was made of acrylic glass material and featured a height-adjustable cover along with designated water entry and drainage points. Four LED lamps were mounted on adjustable side brackets around the arena, allowing control over their height, position, and angle (Fig. S3b). For recording we used the Basler CoaXPress 2.0 boA9344-70cc (Basler, Ahrensburg, Germany) with the GMAX3265 CMOS sensor, capable of delivering 70 frames per second with a 65 MP resolution (9344 × 7000 pixels). It is equipped with a macro lens, the Qioptiq Apo-Rodagon-D 2x 4/75 (Qioptiq/Excelitas, Göttingen, Germany). The camera was installed on motorised linear rails above the arena for X-Y translation, and its position was controlled using a joystick to keep the area of interest within the frame. The camera could be moved on the z-axis through a fine translation stage (Fig. S4). Acquiring videos at the camera’s native full-sensor resolution (9344 × 7000 px; 65.4 MP) exceeded our data-throughput capacity: software compression was too slow and hardware compression introduced visible artifacts. We therefore recorded a square ROI covering ∼25% of the sensor (3968 × 3968 px at 20 fps), corresponding to a 400 × 400 µm field of view (≈ 9.9 px/µm; 0.10 µm/px). This adjustment reduced the total image size but did not alter spatial sampling density, as the physical magnification and pixel size remained identical. This adjustment was not considered a significant loss of information, as the majority of the full frame was out of focus, due to the non-flat curvature of the animal’s mantle.

### Sedation

We used ethanol as a sedative during imaging. To achieve a state of sedation, the animal was gradually exposed to ethanol dissolved in artificial seawater, with the concentration incrementally increased up to 1.5%. The procedure was carried out in a large, dark bucket to reduce visual stimulation. This controlled approach minimised stress on the animal while ensuring effective sedation.

Ethanol is widely used in cephalopod research for its accessibility and reversibility (Andrews et al., 2013; Gleadall, 2013), but could in principle suppress neural activity or alter muscle responsiveness. In our dataset, however, PCA revealed near-identical component counts in sedated and awake head-fixed animals, and motor-unit size (in chromatophores) remained unchanged. Thus, ethanol neither artificially inflated nor suppressed the apparent number of motor subunits, validating its use as a non-invasive means to stabilize animals during high-speed recordings.

### Marking procedure

Because the entire mantle of *Sepia* could not fit under the camera view, it was essential to consistently image the same skin region across recording sessions. To solve this issue we used Visible Implant Elastomer Tags (Northwest Marine Technology, Inc. n.d.), a small and practical silicon-based tagging method. To apply the tag, animals were anesthetised with 1.2 % ethanol and a drop of silicon-base tag was injected through a syringe in the skin (Fig. S5). The silicon was coupled with a curing agent and the tag solidified within 24 hours. This type of tag has been shown to be very effective in cephalopods (Barry et al., 2011; Brewer et al., 2012; Zeeh et al., 2009). In our case, the skin of both *Euprymna* and *Sepia* retained the tag for the entire lifespan and veterinarian analyses showed that the tagging did not cause any form of infection. A blue light lamp attached to the camera helped us find the tag by making it fluoresce. We then always filmed the area of skin around the same tag.

### Head fixation

To enable long filming sessions, we utilised head fixation, allowing us to film a single area continuously with only minimal camera adjustments during recording (Fig. S6). Animals were transferred to a bucket containing 1.7% ethanol in ASW, and transferred to a shallow tank (1.5% ethanol in ASW, continuously bubbled with oxygen) at the onset of surgical plane anesthesia. The dorsal head was raised 1 cm above the water line by a silicone-coated head rest, and the gills were superfused with tank water at a rate of 60 ml/min via soft silicone tubes inserted bilaterally into the mantle cavity. Breathing rate was maintained by observation of the collar musculature and changes in ethanol concentration. Lidocaine (2%, 0.2ml) was injected subcutaneously before exposing the dorsal head cartilage, reapplying lidocaine and removing overlying connective tissue. A thin layer of Vetbond (3M) was then applied to the dorsal head cartilage before attaching a custom 3D-printed head plate (biocompatible resin) with dental cement (Venus, Kulzer) before sealing the wound with surgical glue and transferring the animal to a recovery tank.

### Filming sessions

*E. berryi*: 14 animals were filmed. These recordings were used to establish experimental conditions and to develop and refine the analysis pipeline. To maximize spontaneous chromatophore activity, individuals were lightly sedated with 1–1.5% ethanol, which also induced slight chromatophore contraction. Expansion was triggered by cyclically varying the ambient light intensity with a custom, remotely controlled LED ring positioned above the arena, taking advantage of the light-dependent dynamics of chromatophores (Zylinski et al., 2011). Filming sessions varied in duration depending on the experimental condition. Sedated animals were recorded for up to 20 minutes before being returned to normal seawater for recovery, whereas non-sedated individuals could be filmed for up to 1 hour with continuous seawater circulation. Whenever possible, the same individuals were recorded every three days over the course of development, spanning a total period of approximately two months. The results presented here are based on recordings from a lightly sedated individual exposed to cyclic changes in ambient light intensity. This condition produced the most stable and interpretable chromatophore dynamics, minimizing motion artifacts while maintaining spontaneous activity.

*S. officinalis*: Over the course of the study, 12 individuals ranging in age from 5 days to 10 months post-hatching were filmed under various experimental conditions. A range of experimental configurations was explored to optimize chromatophore visibility and recording stability, including evoked camouflage using printed or e-ink patterns presented beneath a transparent tank, light ethanol sedation (up to 1.5 %), and head fixation. In selected individuals, the same skin region was imaged at three-day intervals for a maximum duration of six months. The dataset presented here was obtained from a head-fixed individual filmed on days 1, 2, and 6 post-surgery, both with and without light ethanol sedation. Head fixation provided stable, high-quality recordings with precise spatial correspondence between frames. Non-sedated sessions lasted up to 2 hours, while sedated sessions were limited to 20-minute bouts before recovery in normal seawater. Comparable results could be obtained from lightly sedated animals when movement was compensated by a tracking camera or other gentle immobilization.

### Computational analyses

Chromatophore activity was quantified using the CHROMAS software package (Ukrow, Renard et al., 2025), which processes video data of cephalopod skin to identify individual chromatophores and extract their spatiotemporal dynamics. CHROMAS provides per-chromatophore metrics such as surface area, epicentre coordinates, and anisotropic deformation. The resulting dataset provided spatiotemporal maps of chromatophore activity, forming the basis for subsequent neural and biomechanical analyses. CHROMAS was used as described in Ukrow, Renard et al., 2025, with the following parameter and model choices.

For chunking, which filters out frames likely to impair analysis, we chose the difference-of-Gaussians (DoG) focus metric with kernel sizes k1=11, k2=5, and standard deviations s1=2 and s2=1 pixels.

To generate training data, manually annotated images (3968×3968 pixels) were divided into 512×512 pixel patches. Of these, 90% were randomly selected for training and the remaining 10% were held out for testing. To improve model generalisation, training data augmentation followed Ukrow, Renard et al., 2025 with the following parameters: spatial translations (±20%), scaling (±20%), rotation (±30°), RGB channel shifts (±25), brightness variation (±10%), contrast alteration (±30%), perspective warping (scale 0.05–0.5), and vertical/horizontal flipping. Each transformation was applied independently with a probability of 25%, except flipping, which was applied with a 50% probability.

Segmentation used the neural-network option of CHROMAS using a Fully Convolutional Network (FCN) architecture with a ResNet-50 backbone (Long et al., 2015). Training was performed for 500 epochs with validation every 5 epochs, using a composite loss function combining Dice loss and cross-entropy loss in equal proportion (50:50). Optimisation used the Adam optimiser with an initial learning rate of 0.001. A *ReduceLROnPlateau* scheduler (mode: ‘min’, factor: 0.1, patience: 50 epochs) adaptively reduced the learning rate when validation performance plateaued. The model with the lowest recorded validation error was retained for analysis.

For registration, 900 randomly selected tracking points were initialised on a 100×100 pixel grid. Displacement vectors exceeding 10 pixels were excluded. The interpolation parameter of the moving-least-squares algorithm was set to α = 3.0.

For stitching, we applied the ellipse-fit option for *Euprymna berryi* and the manual option for *Sepia officinalis* as detailed in Ukrow & Renard, 2025. The interpolation parameter of the moving-least-squares algorithm was set to α = 3.0.

All other parameters were set to their default values. To minimize analysis bias, the pipeline was fully automated and applied using identical parameters across all conditions and datasets.

### Chromatophore anisotropic deformation tracking

Chromatophore anisotropic deformation tracking provides a detailed characterisation of the directional and irregular expansion patterns of chromatophores. This analysis begins with a stabilisation step that compensates for both global and local deformations of the skin. To achieve this, the CHROMAS pipeline identifies stable points on the skin, referred to as motion markers, which correspond to chromatophores that remain small and constant in size across frames. These motion markers are used to derive skin deformation over time, creating a stabilised coordinate system that eliminates artifacts caused by skin movement. Each chromatophore is divided into 36 radial slices to enable precise monitoring of its anisotropic deformations. This number of slices, determined based on the Nyquist–Shannon theorem (Shannon, 1948) and histological analysis (Fig. S7), was set to 36, corresponding to twice the upper bound of histologically confirmed muscle fibers. Using a higher resolution would mainly amplify small segmentation imperfections, the primary source of error. The epicenter of each chromatophore is used as the origin for radial slicing and is calculated from the chromatophore’s center of mass in its most contracted state. This epicenter remains stable across frames due to its relative positioning within the local coordinate system of motion markers. Slice areas are calculated by averaging distances from the epicenter to the chromatophore’s edges, followed by squaring and scaling. This method ensures accuracy even in the presence of pixel discretization errors. Orientation of slices is maintained consistently across frames using the local coordinates derived from motion markers, allowing for the reliable tracking of individual slices over time.

### Expansion and contraction speeds

We analyzed the expansion and contraction dynamics of chromatophores by tracking 36 radial slices for each of ten particularly active chromatophores. For each recording, area values were reshaped into a matrix of dimensions “number of frames × 36 slices” and converted to radii, assuming equal-angle sectors, using the relation r = sqrt(36 × area / π). Peak-centered events were defined for each slice after applying a light Savitzky–Golay smoothing (window size 5, polynomial order 2). Local maxima were detected on the smoothed radius traces using the *scipy.signal.find_peaks* function with a prominence threshold of 1.5. For each detected peak (at frame p), the onset of expansion was defined as the frame immediately before the first positive slope encountered when scanning left from the peak, and the end of contraction as the last frame with a negative slope when scanning right. Slopes were computed from the first difference of the radius signal, with a small threshold (epsilon = 1e-5) used to avoid spurious sign flips due to numerical jitter.

For each event, we calculated expansion amplitude (*A_exp_ = r_p_* − *r_start_*), duration (*D_exp_* = p − start), and mean speed (*S_exp_ = A_exp_ / D_exp_*), as well as the corresponding contraction amplitude, duration, and speed (*A_con_ = r_p_* − r*_end_*, *D_con_* = end − p, *S_con_ = A_con_ / D_con_*). To suppress small fluctuations that likely represented noise, we excluded events with an expansion amplitude smaller than 15% of the full amplitude range. Events were then grouped by expansion amplitude into three categories based on percentile tails (tail width = 33%): “small” (lowest 33%), “center” (middle 33–67%), and “large” (highest 33%). Within each group, we applied Tukey’s interquartile range (IQR) method to remove outliers based on both amplitude and speed, keeping only events that fell within [Q1 − k × IQR, Q3 + k × IQR], with k = 1.

Statistical comparisons of expansion versus contraction speeds and durations at the event level were performed using paired t-tests. Statistical significance was reported as follows: *p* < 0.05 (\**), p < 0.01 (*\*\**), p < 0.001 (*\*\*\*), and *p* < 0.0001 (****). Durations were expressed in frames, equivalent to milliseconds when converted at 20 frames per second (e.g., 50 ms per frame).

### Number and influence of motor neuron per chromatophore

To estimate the number of distinct motor neurons controlling each chromatophore, time series representing the expansion dynamics of radial slices were processed through a dimensionality reduction and source separation pipeline. Each chromatophore’s area trace matrix (frames × slices) was first detrended using *scipy.signal.detrend*. Principal Component Analysis (PCA) was performed using *sklearn.decomposition*. PCA, and the number of meaningful components was estimated via the “elbow method”, with the elbow point identified using the *kneed.KneeLocator* package (Satopaa, 2011). This number was then used as the n_components parameter for Independent Component Analysis (ICA), performed using *sklearn.decomposition.FastICA*. Initial ICA analyses showed that single components sometimes spanned both poles of a chromatophore, due to anticorrelated activity between opposing sides caused by the elastic mechanics of the pigment sacculus. To focus only on *active* motor signals, we filtered the data to retain only expansion events, resulting in independent components localized to one side and providing a clearer mapping of motor neuron influence.

The resulting ICA mixing matrix provides the contribution of each independent component to each radial slice. To quantify this influence, the absolute values of the mixing matrix were normalised per slice, yielding a percentage influence profile across components. These influence profiles were visualised using polar plots to assess spatial distribution. The number of components (ICs) retained per chromatophore serves as an estimate of the number of distinct motor neuron inputs driving that chromatophore’s expansion.

### Number of chromatophores per motor unit

Motor units were inferred by clustering independent components (ICs) extracted from chromatophore activity traces using the FastICA algorithm (*sklearn.decomposition.FastICA*). To group ICs with similar temporal profiles, we applied Affinity Propagation (*sklearn.cluster.AffinityPropagation*) using pairwise Pearson correlation as the similarity metric. Each resulting cluster was interpreted as a putative motor unit, consisting of ICs presumed to originate from the same motor neuron. Chromatophores were then assigned to motor units based on their strongest IC loading. Motor unit size was defined as the number of unique chromatophores associated with each cluster. All steps were implemented in Python (RRID:SCR_008394) using *NumPy* (Harris et al., 2020) (RRID:SCR_008633), *pandas* (McKinney, 2010) (RRID:SCR_018214), and *scikit-learn* (Pedregosa et al., 2011) (RRID:SCR_002577).

### Motor unit shape and structure

To characterize chromatophore clusters, we developed a Python script using *pandas* and *NumPy* for data handling, *SciPy* for distance calculations and minimum spanning tree construction, and *matplotlib* for visualization (Hunter, 2007) (RRID:SCR_008624). To assess the spatial distribution of chromatophores quantitatively, we computed nearest-neighbor distances (NND) using Python. Epicenter coordinates [epicenter_x, epicenter_y] were extracted from our dataset. For the global analysis, we computed pairwise Euclidean distances between all chromatophore epicenters and recorded the minimum distance to identify each chromatophore’s nearest neighbor. For the within-cluster analysis, we grouped chromatophores by cluster label and repeated the NND computation within each group. The output metrics included the mean NND, standard deviation, and coefficient of variation (CV), which were used to describe both overall spatial density and intra-cluster compactness. Results were visualised by plotting chromatophore positions and connecting each to its nearest neighbor with a line segment. In addition to nearest-neighbor distances (NND), we calculated furthest-neighbor distances (FND) and convex hull areas (via *scipy.spatial.ConvexHull*) to estimate cluster extent. From these, density metrics were derived by relating chromatophore counts to convex hull area, and Pearson correlation coefficients (*scipy.stats*) were used to assess relationships between area, density, and number of chromatophores.

To visualize the putative motor units’ ramifications, chromatophores were grouped by cluster label and plotted according to their epicenter coordinates, with multi-labeled chromatophores displayed as multicolored wedges, and internal cluster geometry was quantified by computing pairwise Euclidean distances and reducing them to a minimum spanning tree. This framework enabled both visualization and quantitative comparison of cluster compactness, extent, and density.

### Second order innervation

To assess second-order innervation, we first extracted all combinations of chromatophore identifiers and their corresponding cluster labels from the dataset, yielding a unique set of cluster memberships. A pairwise co-occurrence count was computed by iterating over all clusters and recording how often each pair of chromatophores appeared together; we then counted how many pairs occurred together in two or more clusters. For the null model, we implemented a constrained permutation in which chromatophore IDs were randomly reassigned across clusters while preserving the number of unique IDs per cluster. This procedure maintains the original cluster sizes but not the per-chromatophore memberships. Across 1,000 simulations, we repeated the pairwise counting procedure and recorded the number of pairings that reached the second-order threshold. This produced a distribution of null values, against which the real count was compared using a two-tailed z-test to determine statistical significance.

### Electrophysiology

#### Animals

Electrophysiological experiments were carried out in Beijing, China. At the time of the experiments, there were no specific national regulations governing the use of cephalopods in research in China. Nevertheless, all procedures performed at Peking University were designed to comply, as far as possible, with the principles of the European Directive 2010/63/EU and FELASA recommendations.

For electrophysiological experiments, bobtail squids *Euprymna berryi* were either hatched from eggs laid in house or purchased as eggs and naturally hatched in the lab. Both adult squids and eggs were sourced from Guangdong and Hainan provinces in China. *E. berryi* were reared in an artificial seawater (ASW) system around 24 □, with a salinity of 28 ppt and a pH of 8.3. Water quality was monitored continuously and was tested regularly. The ASW filtration system consists of filter floss, filter bags, biofilter (for nitrifying bacteria), a protein skimmer, and UV-lamp. Room light was set as a 12h-12h light-dark cycle with gradual on- and off-sets at 12:00 and 24:00.

#### Brain and skin preparation

Juvenile *E. berryi* ranging from 5-10 mm in body length were used for electrophysiological experiments. Animals were first anesthetized by 2% ethanol in ASW, and then transferred into an acrylic chamber with a sloped silicone rubber base, filled with oxygenized calcium-free saline (460 mM NaCl, 10 mM KCl, 10 mM glucose, 10 mM HEPES, 55 mM MgCl_2_, and with a pH of 7.4.) at room temperature. The arms, beak, and both eyes were removed. The mantle was opened from the ventral side and all visceral organs were removed. The skin of the head was removed to expose the posterior sub-esophageal mass. The posterior chromatophore lobe was de-sheathed gently with fine forceps. The mantle was pinned on a 70° sloped silicone rubber plane, while the brain was placed on a flat silicone rubber base. The mantle and the brain were connected solely through pallial nerves. The solution was then replaced by oxygenated (100% O2) calcium-containing saline (460 mM NaCl, 10 mM KCl, 10 mM glucose, 10 mM HEPES, 55 mM MgCl_2_, 11 mM CaCl_2_, and with a pH of 7.4).

#### Electrophysiological recording and imaging setup

The patch clamp recording was performed under an upright fluorescence microscope (BX51W1, Olympus, Tokyo, Japan) with a 20 water-immersion objective (UMPlanFI 20, Olympus, Tokyo, Japan). The brain was visualized using a microscope camera (Moment, Teledyne Vision Solutions, Thousand Oaks, California, USA). The mantle chromatophore was recorded using a camera (a2A4096-30ucBAS, Basler, Ahrensburg, Germany) with a manual lens (MLM-3X-MP, Computar, Tokyo, Japan) at 30 fps. The glass electrodes (∼8 M) were made from borosilicate glass (BF150-86-10, WPI, Sarasota, Florida, USA) by a puller (P-1000, Sutter Instrument, Novato, California, USA). The pipettes were filled with internal solution (450 mM K-gluconate, 10 mM NaCl, 4 mM MgCl_2_, 3 mM EGTA, 20 mM HEPES, 2 mM Mg-ATP, 0.2 mM Na-GTP, 2% neurobiotin, and with pH of 7.4). Recordings were performed in current clamp mode with a HEKA amplifier (ESC100-USB, Lambrecht, Germany). For intracellular stimulation, step or constant currents were injected repetitively at 0.5 Hz. The onset of the stimulation train simultaneously triggered the camera to record chromatophore activity on the mantle skin.

## Data Availability

The image analysis software CHROMAS is distributed via the pypi package index (https://pypi.org/project/chromas/) and is publicly released on GitLab (https://doi.org/10.17617/1.pa38-mh49) under the 3-Clause BSD License. The documentation is hosted on GitLab.

Code and example datasets used for the biological analyses reported in this study are provided in the “motor_units” folder at https://public.brain.mpg.de/ (under the project directory named after this paper). Electrophysiology videos and trace data are provided in a separate folder named “ephys”.

## Acknowledgments

We thank F. Bayer for assistance in building the experimental setup; F. Kretschmer for optimizing the camera control, recording software and Git workflows; F. Vollrath and S. Junek for help with image acquisition; P. Musset for help with high performance computing; L. Jürgens, S. Schwind, L. E. Reyes de Frey, D. Burgard, M. Landler, M. Minde, S. Kranz, P. Dominiczak, N. K. Vogt, G. Wexel, S. Dizdarevic and E. Northrup for animal care; and T. Woo and other members of the Laurent laboratory for constructive exchanges. We thank Alice Perenzin and Antje Berken for grant management and scientific coordination. This research was funded by the Max Planck Society (GL), by the LOEWE Schwerpunkt CMMS (State of Hesse) (GL), by the Louis Jeantet Foundation (GL), by the European Union (ERC grant CAMOUFLAGE, 10114150) (GL), by National Natural Science Foundation of China (NSFC grant 32371215) (XL), State Key Laboratory of Membrane Biology, Peking University (XL); and Peking-Tsinghua Center for Life Sciences (XL).

---

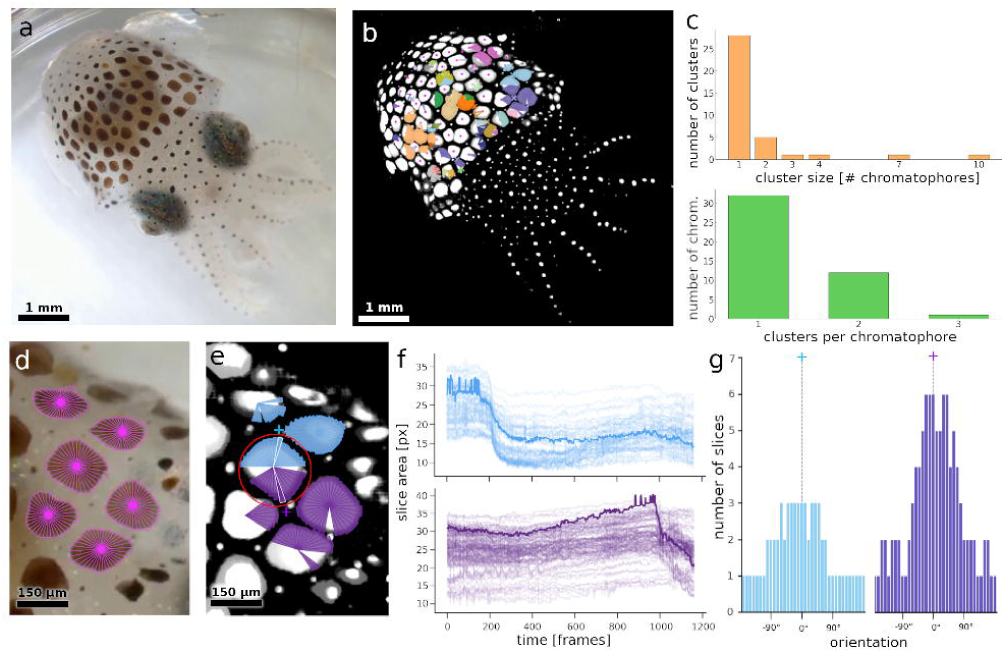

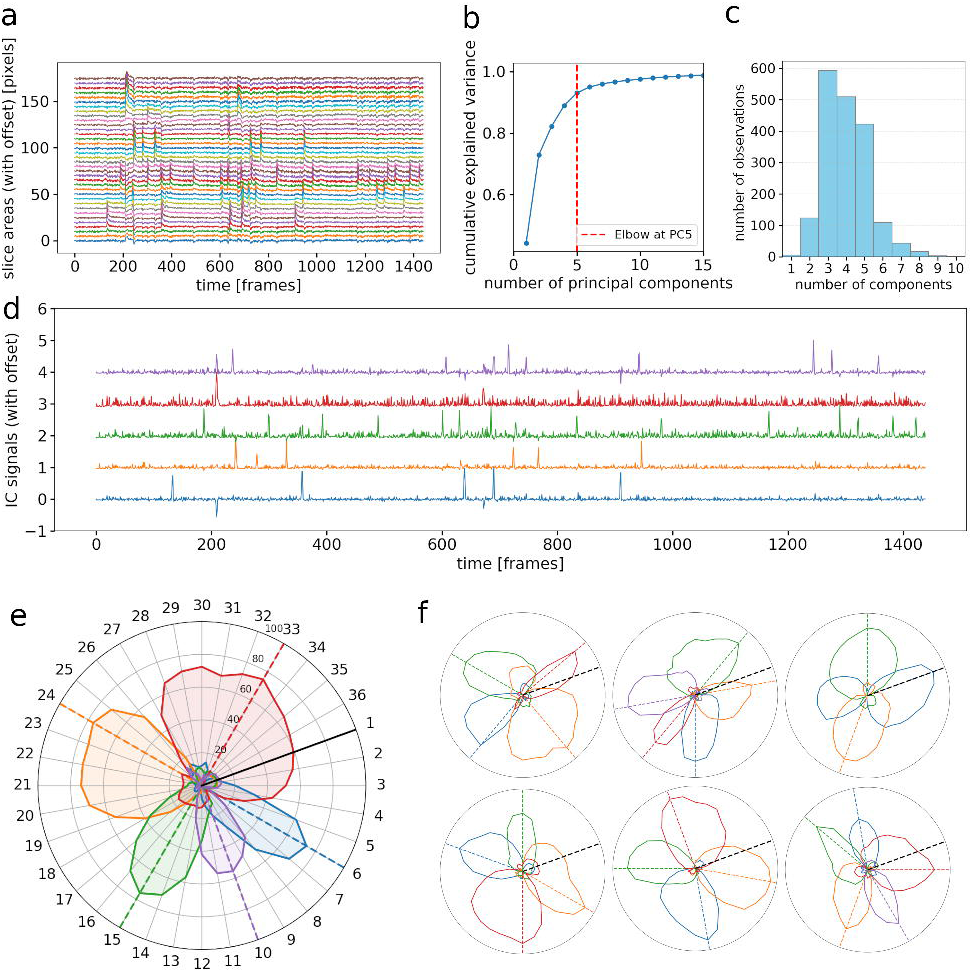

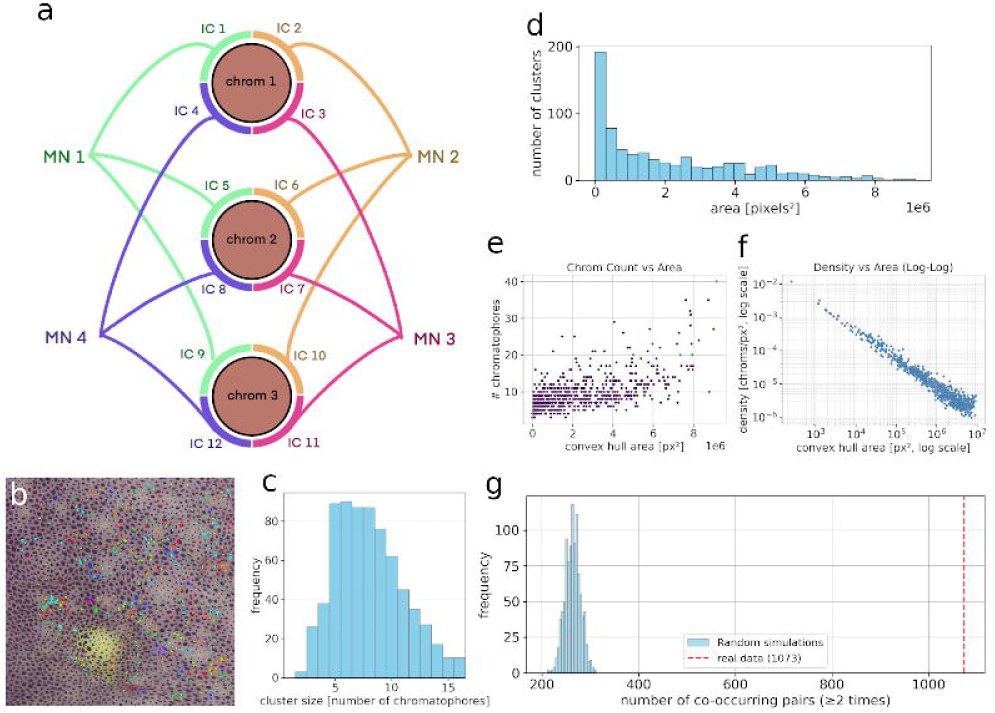

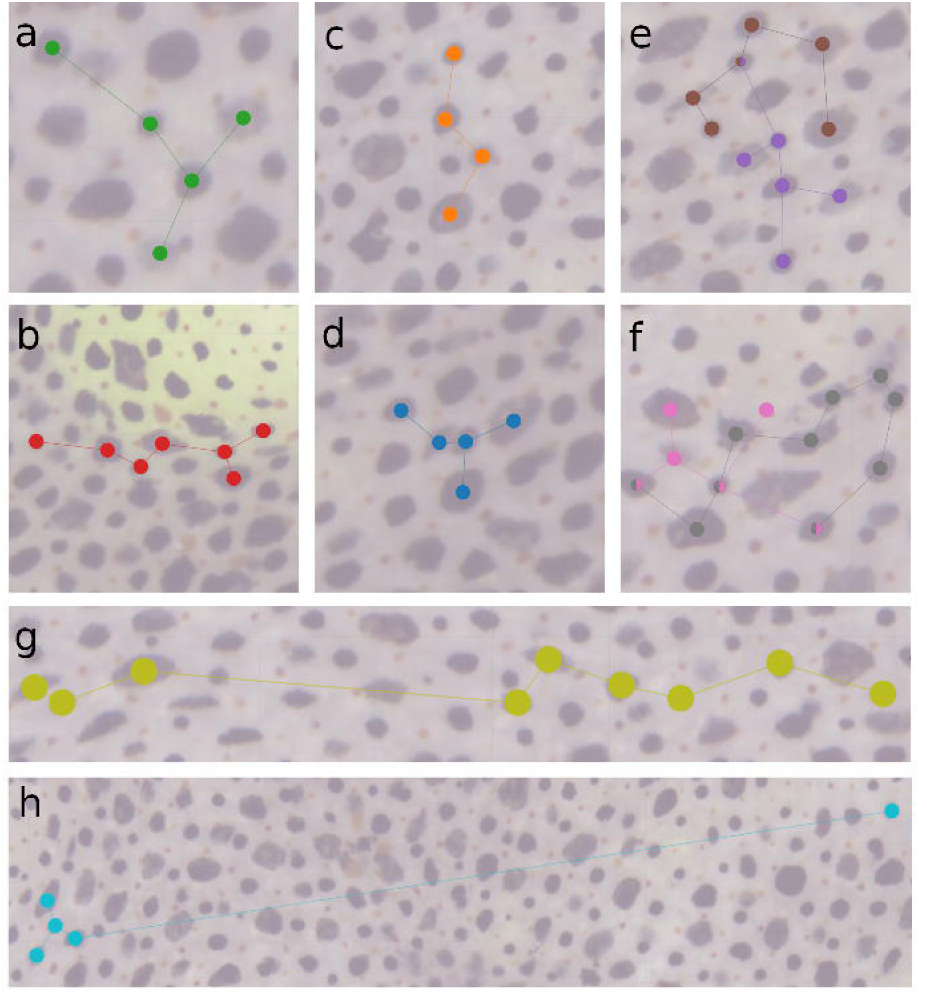

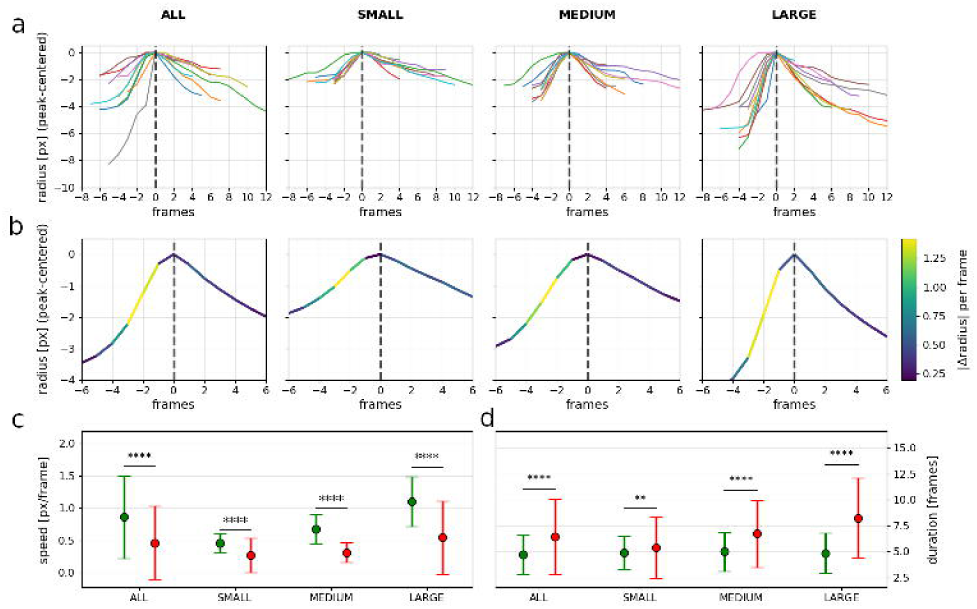

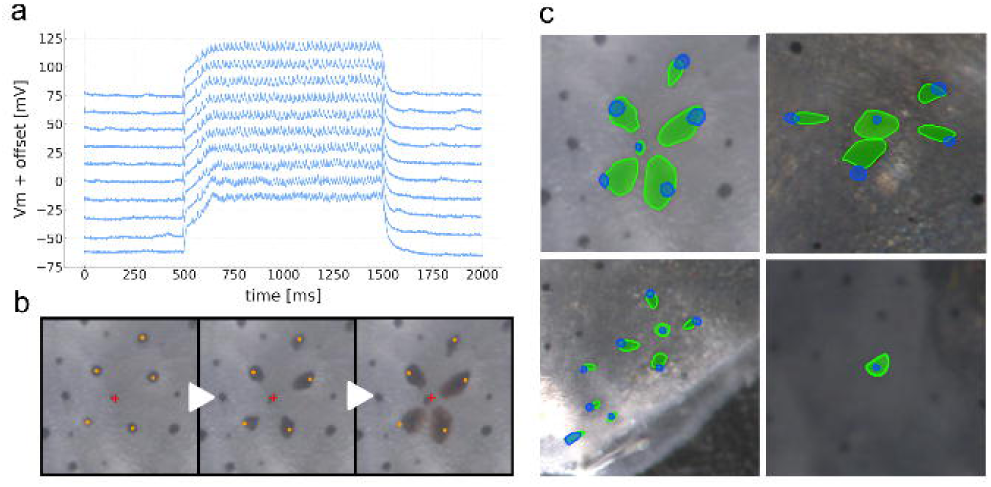

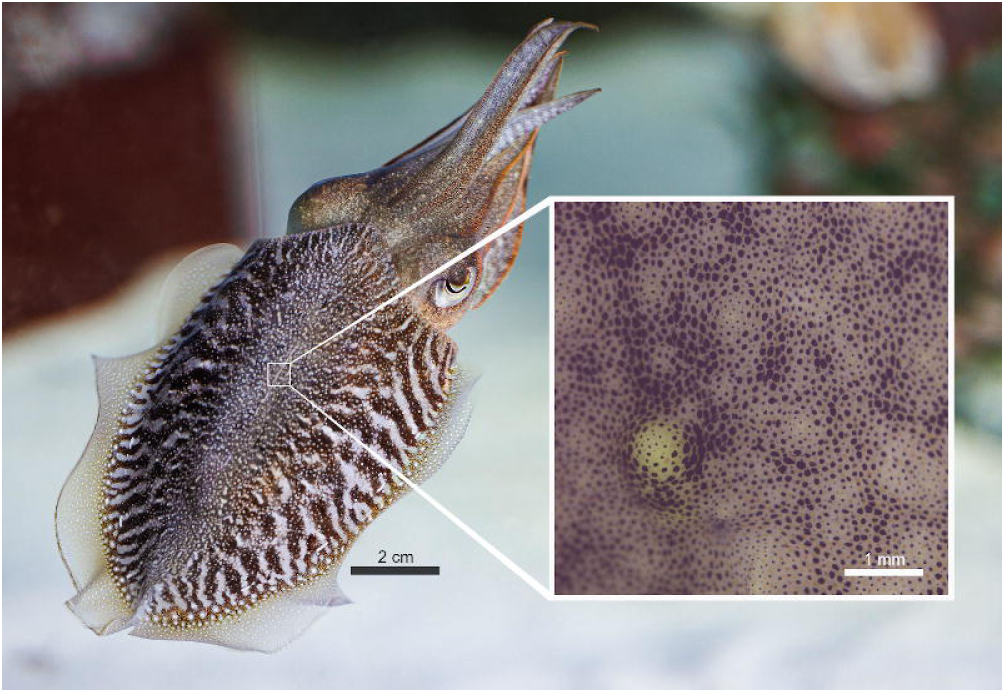

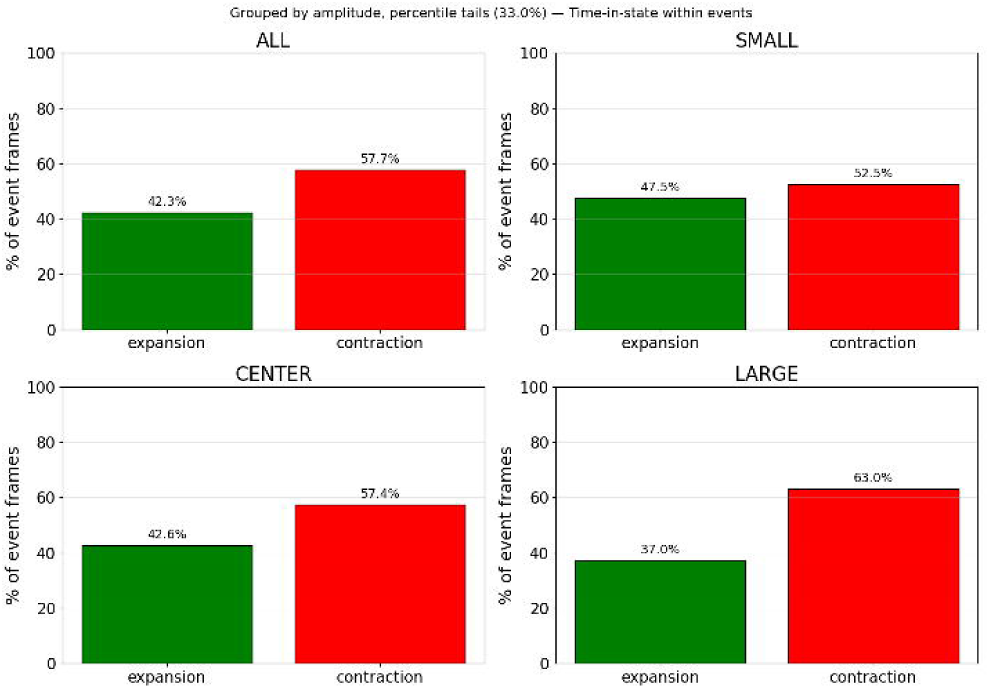

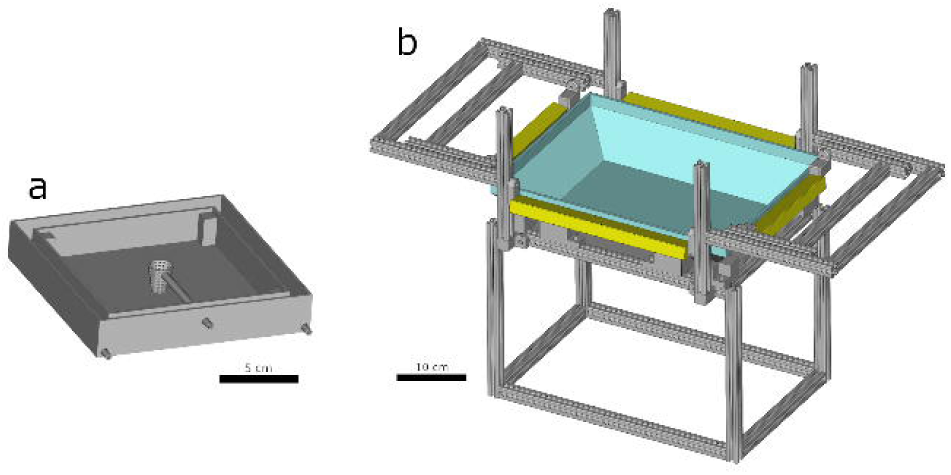

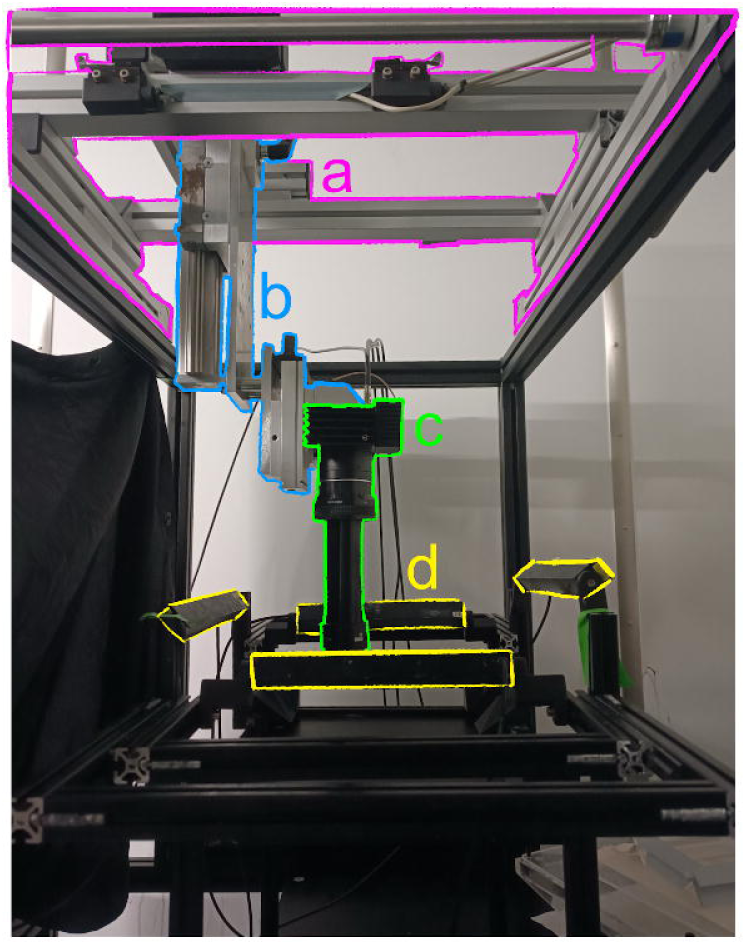

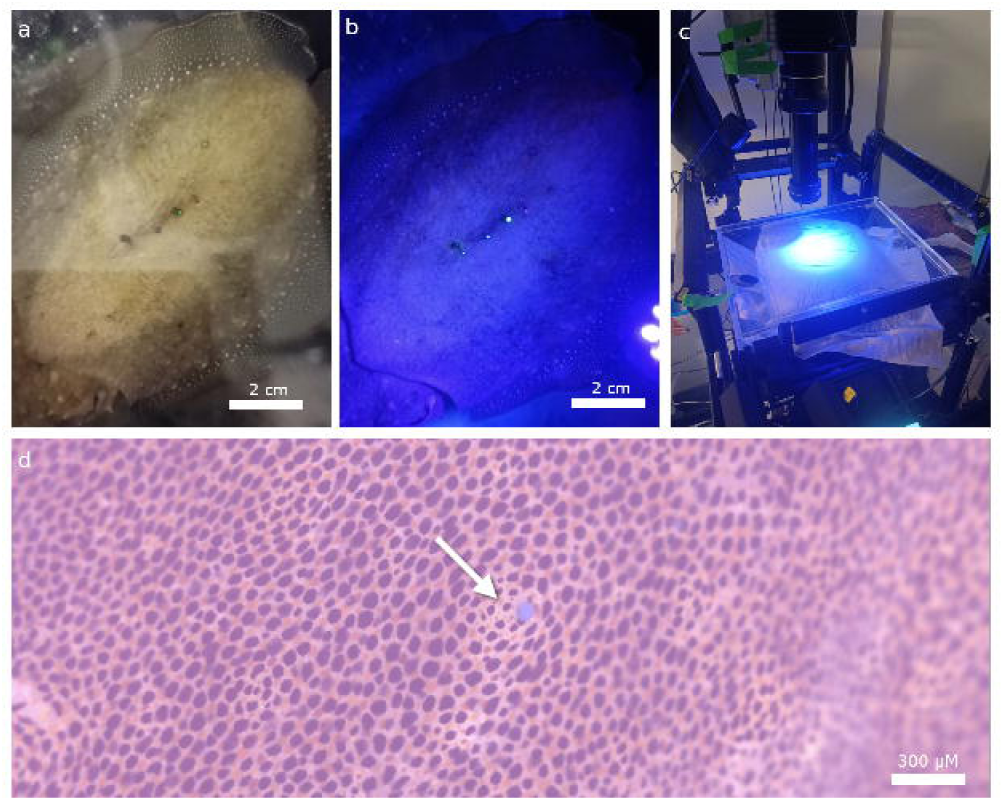

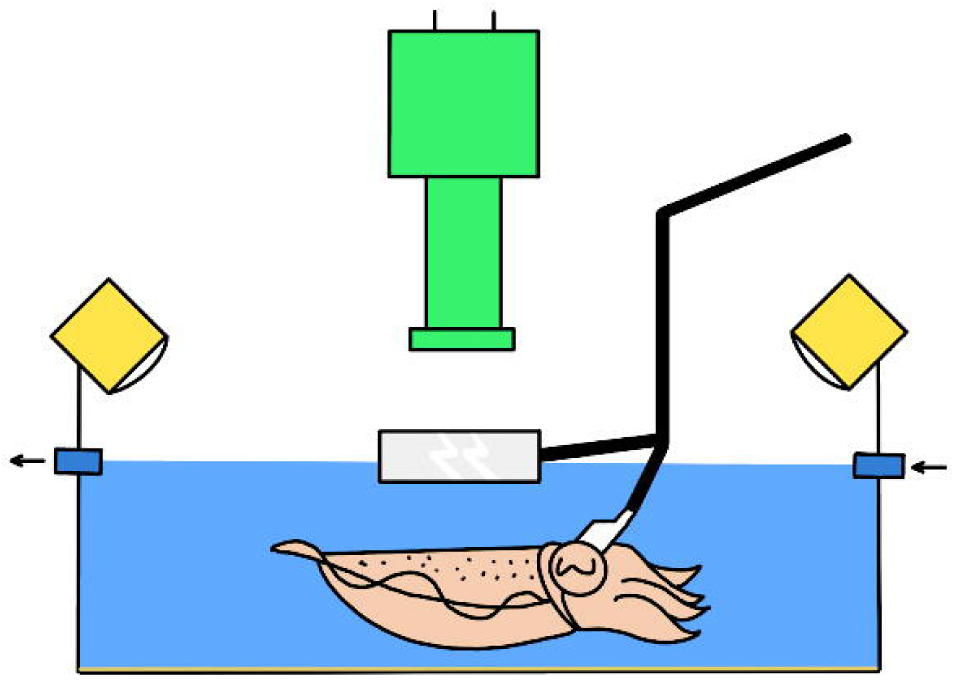

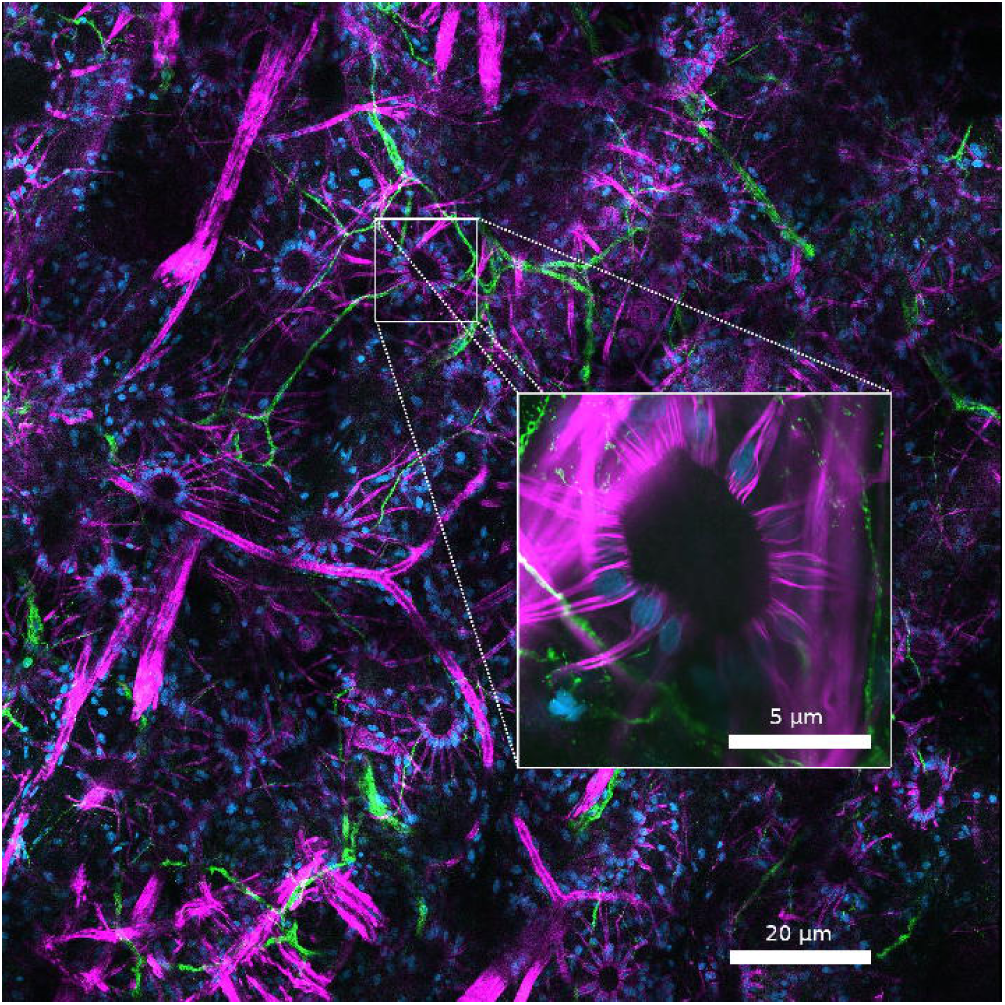

